# I2Bot: an open-source tool for multi-modal and embodied simulation of insect navigation

**DOI:** 10.1101/2024.07.11.603012

**Authors:** Xuelong Sun, Michael Mangan, Jigen Peng, Shigang Yue

**Author notes:** **Author for correspondence:** Shigang Yue; Jigen Peng.

## Abstract

Achieving a comprehensive understanding of animal intelligence demands an integrative approach that acknowledges the interplay between an organism’s brain, body, and environment. Insects like ants, despite their limited computational resources, demonstrate remarkable abilities in navigation. Existing computational models often fall short in faithfully replicating the morphology of real insects and their interactions with the environment, hindering validation and practical application in robotics. To address these gaps, we present I2Bot, a novel simulation tool based on the morphological characteristics of desert ants. This tool empowers robotic models with dynamic sensory capabilities, realistic modelling of insect morphology, physical dynamics, and sensory capacity. By integrating gait controllers and computational models into I2Bot, we have implemented classical embodied navigation behaviours and revealed some fundamental navigation principles. By open-sourcing I2Bot, we aim to accelerate the understanding of insect intelligence and foster advances in the development of autonomous robotic systems.

## 1. Introduction

Unravelling the underlying mechanisms that give rise to animal’s intelligent behaviours represents a formidable challenge, one that demands an integrative approach that transcends conventional boundaries. Central to this endeavor is the recognition of the dynamic interplay between an organism’s brain, body, and environment [1, 2] – a concept central to the biorobotics approach [3–5]. By bridging the gap between biology and robotics, biorobotics offers a unique opportunity to gain insights into the mechanisms underlying intelligent behaviour [3, 6] while simultaneously inspiring the development of more advanced robotic systems [7–9].

Insect navigation stands as a remarkable example of intelligent behaviour, offering a fertile ground for exploration [10]. The intelligence evidenced by insect navigation [11, 12] which includes both adaptive behaviours that changes dynamically as the environment/stimuli changes [13–15] and cognitive behaviours involving complex decision-making and information integration [16–18]. Despite possessing compact neural architectures with limited computational resources, insects demonstrate an astonishing capacity to navigate complex environments [19, 20], integrating multisensory inputs [16, 21], learning from experience [22], and executing adaptive decision-making processes [23–26]. This ability to solve intricate problems with constrained computational power has sparked profound curiosity and a desire to understand the principles that govern these behaviours, with applications possible in AI and robotics [12].

Computational models play important roles in revealing the neural circuits underlying insect navigation [27–29], and some of these models have been verified using mobile robots [28, 30, 31]. However, these robots could not fully recapitulate the real insects’ body due to differences in morphology. Simulations offer significant advantages over real-world robots when studying complex systems, such as the ability to conduct controlled experiments flexibly, modify parameters easily, and rapidly iterate through multiple scenarios [32]. Although neuromechanical simulation tools for *Drosophila* have recently been developed [33–35], there remains a lack of tailored, realistic simulation tools for the systematic and comprehensive validation of insect navigation models, particularly for desert ants, that account for physical constraints.

To address these limitations, by leveraging the user-friendly and open-sourced software - ***Webots*** [36], a widely used robot simulation tools in academia [37–40], we developed a simulation tool that incorporates the morphological characteristics of desert ants, endowing our navigating agent with dynamic vision, olfactory, tactile, and mechanosensory capabilities (see Figure 1). Leveraging this framework, we have implemented simple Forward Kinematics (FK) and Inverse Kinematics (IK) gait controller, enabling the realisation of anatomically constrained computation model for path integration (PI). Additionally, our simulation tool facilitates the integration of vision and olfactory senses, allowing us to explore the sensory-motor closed loop inherent in insect navigation. Through systematic testing, we have uncovered evidence of how body-environment interactions can simplify the design of control models (see Table 1). Compared with other related simulation tools, the proposed open-source tool offers advantages in multimodal sensory capacity, flexibility in constructing 3D worlds, support for multiple programming languages, lower learning costs, and a user-friendly interface (see Table 4 and Section 4 for more detailed discussion).

**Table 1.** Summary of the presented case studies with I2Bot. Text in red highlights the new insights and contributions.

|  | Sensor | Motor | Environment |
| --- | --- | --- | --- |
| path integration | velocity + direction<br>**Bio-plausible neural output and simple gait control facilitate homing**[28, 31] | 6 legs | plain terrain |
| visual beacons | binocular/monocular vision<br>**Embodied sensory input effects the performance of controller**[1, 2] | 6 legs | plain terrain and 3D trees |
| visual compass | panoramic vision<br>**How internal representation of direction could control body orientation**[41, 42] | 6 legs | plain terrain and 3D trees |
| odour trail following | odour concentration (stable)<br>**Embodied sensors simplify the design of the controller**[43] | 6 legs + 2 antenna | optic-simulated odour trail |
| odour plume tracking | odour concentration (fluctuated) + wind direction<br>**First implementation of the embodied plume tracking**[13] | 6 legs | virtual wind and odour puffs |

**Figure 1.**
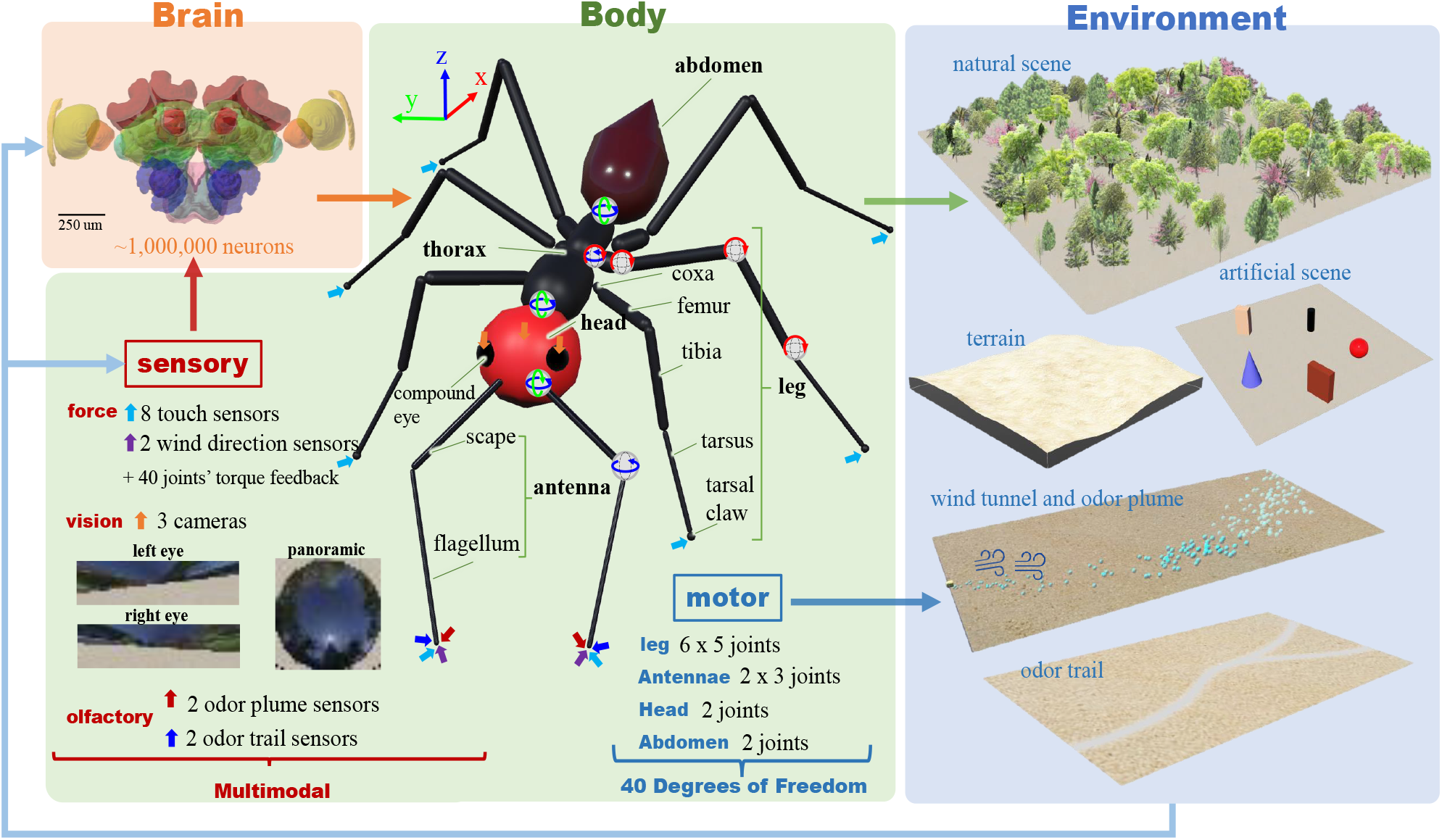
I2Bot overview. The tools aim to integrate simulation of brain, body, and environment, encompassing the sensory-motor loop. Colour-coded arrows indicate the positions of corresponding sensors, while joints are identified by their rotation axes. Note data here is for illustrative purposes, with specific examples shown in subsequent figures.

By open-sourcing our simulation tool, we aim to provide a valuable resource for the research community, accelerating our collective understanding of insect intelligence. Furthermore, we believe that our work holds significant implications for the development of insect-inspired autonomous robots, offering insights that could pave the way for more adaptive and efficient robotic systems [7, 8].

## 2. Results

Example simulated agent/animal and environments are depicted in Figure 1, illustrating the intricate interactions between brain, body, and environment. The neural models can be integrated into the *Controller* component of the Webots’ Robot. The morphology and spatial configurations of individual organs and limbs are reconstructed based on measurements obtained from real desert ants (refer to Section (a) for detailed information). The environments are assembled using Webots’ built-in modules (refer to Section (b) for detailed information).

In this section, we will first introduce the sensory-motor system and the fundamental locomotion capabilities of a simulated desert ant. Following this, as summarised in Table 1, we will present several case studies demonstrating the utilisation of the proposed tool to implement classical control scenarios in a sensory-motor loop fashion. These simulations provide compelling examples of how I2Bot can offer new insights into understanding insect intelligence through the dynamic interaction between brain, body, and environment. In the future we hope for the tool to be extended to include additional sensory modalities, brain models, and body structures of other insects (see Table 5).

Note that force sensing (e.g., mechanosensory, tactile) is not utilized in the current implementation of vision- and olfactory-based behaviours. However, the sensory repertoire can be easily extended to incorporate force sensing data from the antennae and legs in specific scenarios, as these capabilities are already available (see Figure 1).

### (a) Sensors and motors

Insects rely on multimodal sensory information to perceive their surrounding environment and make decisions [44]. Among these modalities, vision, olfaction, and tactile cues are considered crucial for insect navigation. A valuable tool should facilitate easy access to these sensory signals. To address this, the ant robot in I2Bot has been equipped with visual, olfactory, and force sensors. As depicted in Figure 1, two distinct vision sensors—binocular and panoramic—have been incorporated to serve different purposes in visual processing. Similarly, the proposed tool enables simulation of olfactory signals with various spatial-temporal patterns, such as stable odour trails for simulating trail following and dynamic odour plumes for investigating navigation in turbulent olfactory environments. Tactile and mechanosensory input is simulated through force sensors located at the tip of each leg, the tip of each antenna, and torque sensors located in each leg joint (5 per leg), at 3 locations in each antenna, and 2 in the neck and abdomen joints. Finally, wind sensing is simulated by 2 sensors positioning at the tip of the antennae, see Figure 1 for overview.

The locomotion capabilities of insects have long intrigued researchers and have served as inspiration for the design of six-legged robots [4, 8]. To replicate the authentic locomotion of insects, we have defined five degrees of freedom (DoF) for each leg (i.e., front left-FL, middle left-ML, hind left-HL, front right-FR, middle right-MR, hind right-HR). The joints are designated based on the anatomy of body and leg segments. For instance, the joint ThCz refers to the joint connecting the *Thorax* and *Coxa*, rotating along the z-axis. To streamline body movement, the head and abdomen each possess two degrees of freedom, allowing rotation along the y and z axes. Recent studies have emphasised the pivotal role of active olfactory sensing [43, 45, 46] in shaping the navigational behaviours of insects and its application in robotics [47]. Therefore, each antenna of I2Bot is equipped with three degrees of freedom to enable agile movement of the antenna.

### (b) Basic locomotion

Locomotion is the basis of complex manoeuvres that have been observed in insects, thus it is crucial for navigation behaviours. To simplify the locomotion control, we have adapted concepts from robotics and implemented a simple gait control algorithm based on forward and inverse kinematics (FK and IK). Unlike higher level locomotion controllers [48, 49] wherein the gait is assumed fixed or emergent, this approach concerns the movement of each joint (i.e., directly set the angle of each joint) for a given leg movement pattern (i.e., a gait). A kind of lower level method without considering the muscles’ contraction and relaxation [50]. The forward kinematics (FK) method directly assigns the joint angle which determines the position of the leg tip. While the inverse kinematics (IK) computes each joint angle given the spatial position of the end tip of each leg (for more details see Section (c)). That’s to say, in FK, we design the joint angles while in IK, we design the spatial movement of the leg tip and the joint angles are computed through trigonometry. Note that in the current implementation, we only use 3 DoF of each leg (keep angles of the ThCx and the TiTa joint to be constant), which is simpler and more usual for hexapod robots. However, users could in practice apply all the 5 DoF to replicate more biologically realistic locomotion control. In summary, the locomotion models currently included are based on robot control strategies, and could be efficiently upgraded to match models from the biological literature (e.g. [48, 51]) since the joint torque feedback are already accessible.

As depicted in Figure 2 (a) and (b), inverse kinematics-based gait control exhibits greater stability but demands more computational resources. To assess the motion control’s performance, we evaluated its ability to navigate uneven terrain and climb walls. Figure 2 (c) illustrates the comparison of walking on floors with varying degrees of unevenness. Here, we observed faster changes in the z-position of the body with negligible differences in joint torques, indicating the six-legged robot’s inherent capacity to navigate uneven terrain embedded in its physical dynamics. In climbing vertical walls, insects utilise adhesion which we simulated by dynamically applying force to the leg’s ground contact points. Figure 2 (d) presents the climbing performance with varying levels of force applied to the ground support points. Interestingly, larger force doesn’t necessarily translate to better wall-climbing performance. This may be due to the competition between the force required to lift the leg and the force needed to maintain contact with the wall. Different gait types are demonstrated in Figure 2 (g,h), with the tripod gait achieving the fastest body movement speed, while the wave tripod is the slowest. It’s worth noting that other gait controllers such as CPG and feedback models [4, 48, 52, 53] could also be integrated. Here, we present the simple FK-based gait controller to showcase the proposed tool’s feasibility.

**Figure 2.**
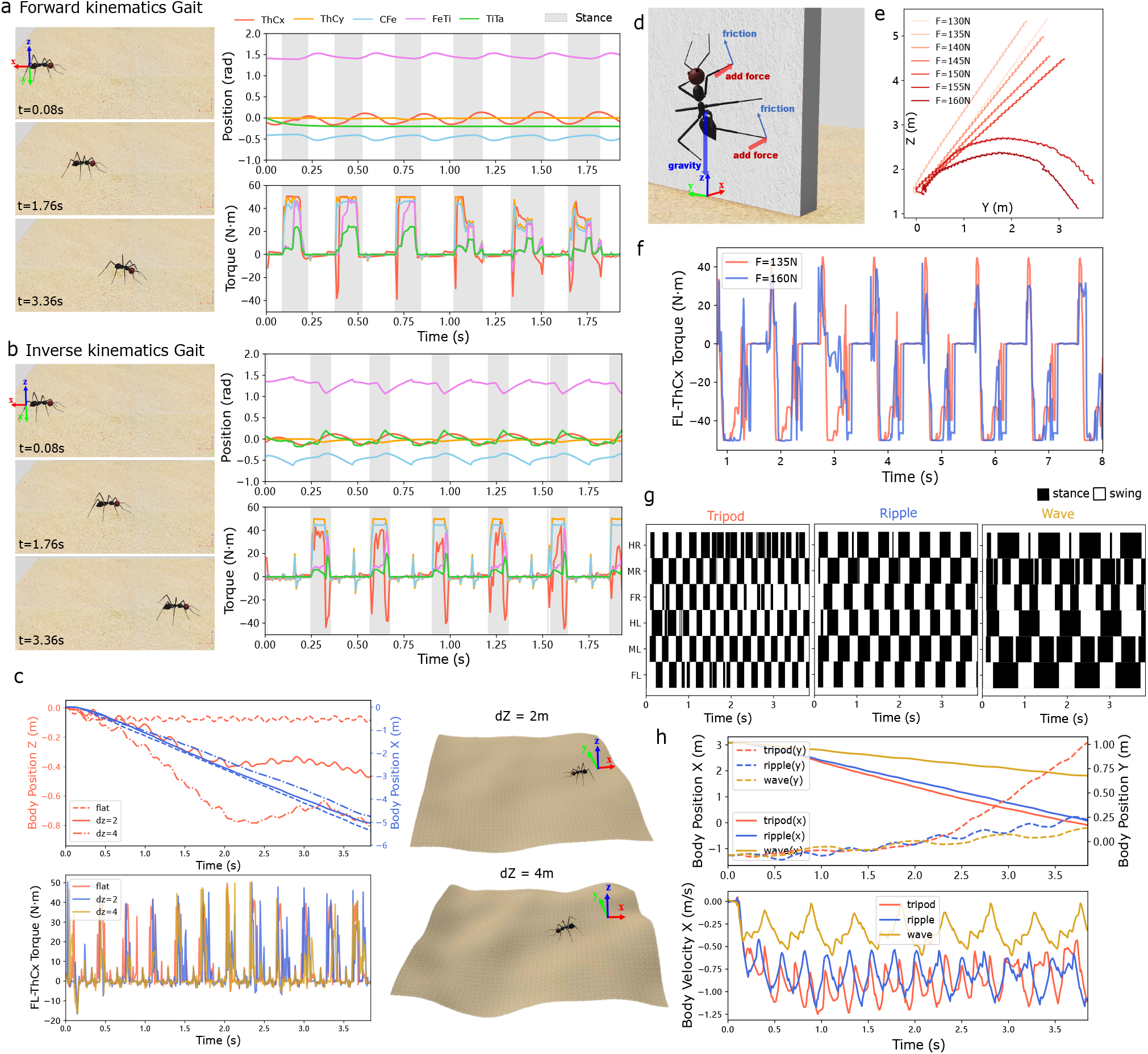
Locomotion and gait control. (a)-(b) Left: snapshots of walking ants at specific times; Right: joint position and torque of the middle right leg based on forward and inverse kinematics. (c) Walking on uneven terrain. (d)-(f) Climbing the wall using adhesion with different force added at the tip of each leg, trajectories are shown in (e) and the torque values are plotted in (f). (g)-(h) Forward kinematics-based implementation of different gait types: tripod, ripple, and wave. Note that when walking on the floor, the robot is initially facing negative x-axis (see the coordinates labelled in Figure 1). All the origins are marked in the figure for clarification.

### (c) Path integration: incorporating locomotion with a bio-constrained neural model

Path integration (PI) stands as a cornerstone of the insect navigation toolkit [54]. Foragers adeptly track the distance and direction to their nest by integrating the series of directions and distances travelled into a *home vector* [19, 55]. Guided by this vector, pointing towards the start position (typically the nest), desert ants can return accurately to their nest even after traversing hundreds of meters (Figure 3a). Recent neuroethological investigations have unveiled the central complex (CX) within the insect brain as pivotal for the computational processes involved in path integration. The CX not only hosts a ring attractor network encoding the animal’s heading direction [56, 57] but also receives optic flow information as a velocity encoding [28, 58].

**Figure 3.**
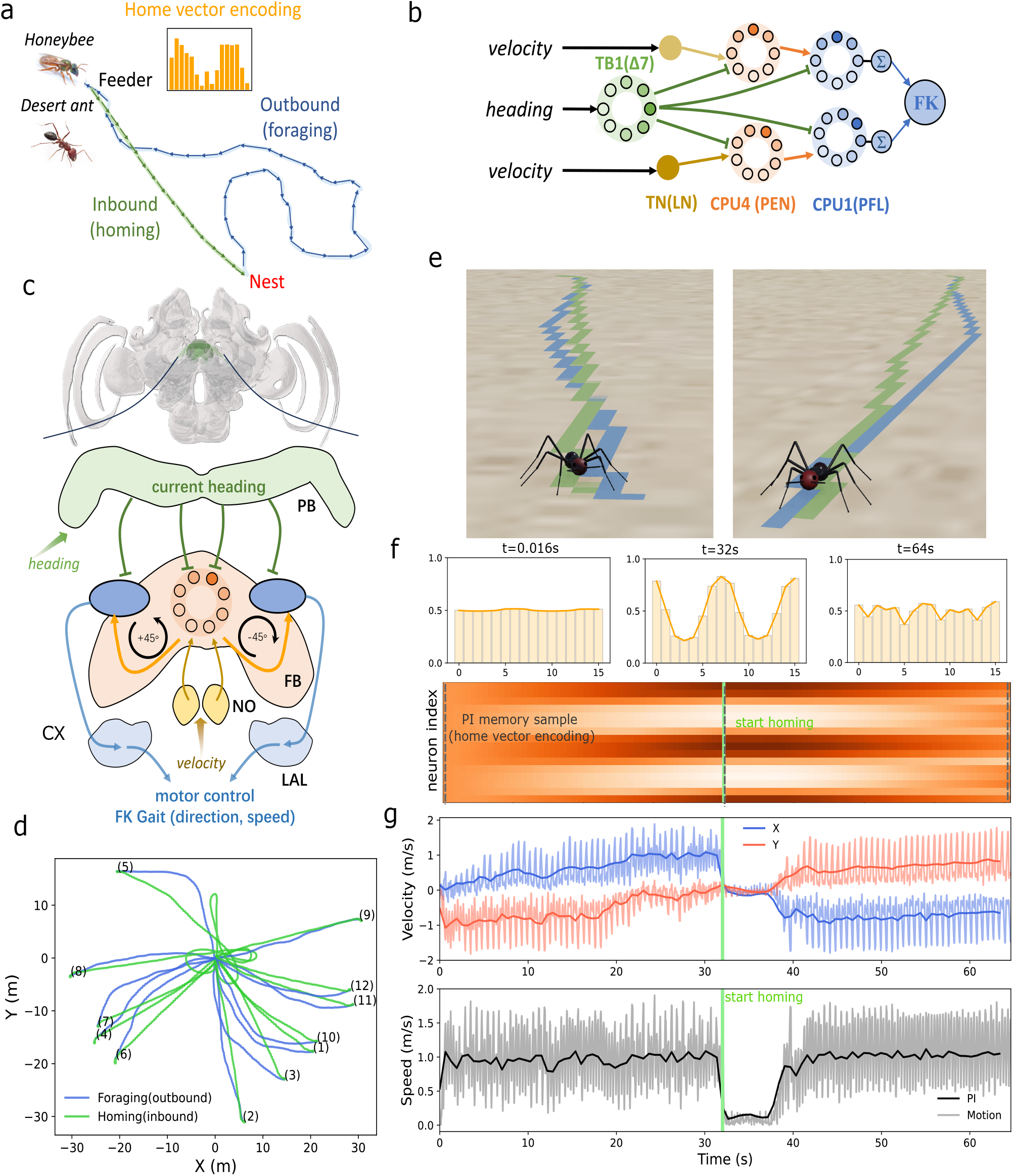
Implementation of path integration model. (a) Schematic diagram of central-foraging insects’ path integration. (b) The neural model of path integration adapted from [28]. The output of the model is linked to the forward kinematics gait controller. (c) Insect brain regions within the central complex related to PI computation. (d) Trajectories of simulated ants conducting foraging trips and homing using the proposed PI model. (e) Snapshots of two typical experiments. Trajectories are drawn using the *Pen* function within Webots. (f) Neural activation of PI memory neurons (CPU4 or FPN neurons) during foraging and homing, with sampled points marked by grey dotted lines. Corresponding activation profiles are placed above. (g) Velocity in the x- and y-axis (top) and speed (bottom) during navigation. Dark solid lines indicate the inputs fed into the PI model, while lighter curves show instant velocity measurements of the walking robot.

To demonstrate the ease of incorporating biologically constrained neural models with locomotion controllers using I2Bot, we implemented the popular insect path integration model proposed by [28]. We utilised its output as tuning factors for the previously described forward kinematics gait controller. Specifically, the neural activation of the left and right PFL/CPU1 neurons from the insect steering circuit modifies the *hip swing* (speed) and *rotation* (direction) of the gait (Figure 3b,c). This marks the first instance of testing this popular neural model in a hexapod robot with physical constraints and ant-like morphology. Notably, the simulated ant can perform curved walking, unlike a recent biorobotics study [31] where the robot could only rotate on the spot. Several simulations were conducted, and we found that this bio-constrained model performed effectively (see Figure 3d,e), suggesting a robust and efficient solution for robot path integration. Figure 3f&g illustrate the dynamic neural activation of the encoded *home vector* and the agent’s velocity during foraging and homing. This exemplifies how neural computation and encoding, coupled with a simple locomotion controller, can generate robust navigation behaviours.

### (d) Vision-guided manoeuvres

Vision serves as one of the primary means for animals to perceive their environment, and insects rely on visual cues for navigation in various ways [59, 60]. In this section, we present scenarios involving monocular, binocular, and widely used panoramic vision to demonstrate vision-motor control in the simulated ant robot.

### (i) Monocular and binocular visual beacons

Research has indicated that insects instinctively use visual landmarks as beacons [61, 62]. To showcase the feasibility of using visual images as input with our proposed tool, we implemented a simple visual beacon algorithm (see Figure 4a,b) based on the biologically plausible steering circuit used in neural models of the insect brain [28, 29, 58]. To compare the effects of different types of visual input on motor output, we employed monocular (i.e., solely left or right image) and binocular images as inputs to the visual beacon model. The results are illustrated in Figure 4d, where the simulated ant with binocular image inputs outperforms the others in terms of task completion time. Eye occlusion resulted in ipsilateral motor bias, where agents with the left eye occluded (receiving only right eye information) approached the landmark with left-biased trajectories (red curves in Figure 4c). This illustrates how embodied sensory input influences navigation performance guided by the same neural controller, providing partial insights into the advantages of binocular vision.

**Figure 4.**
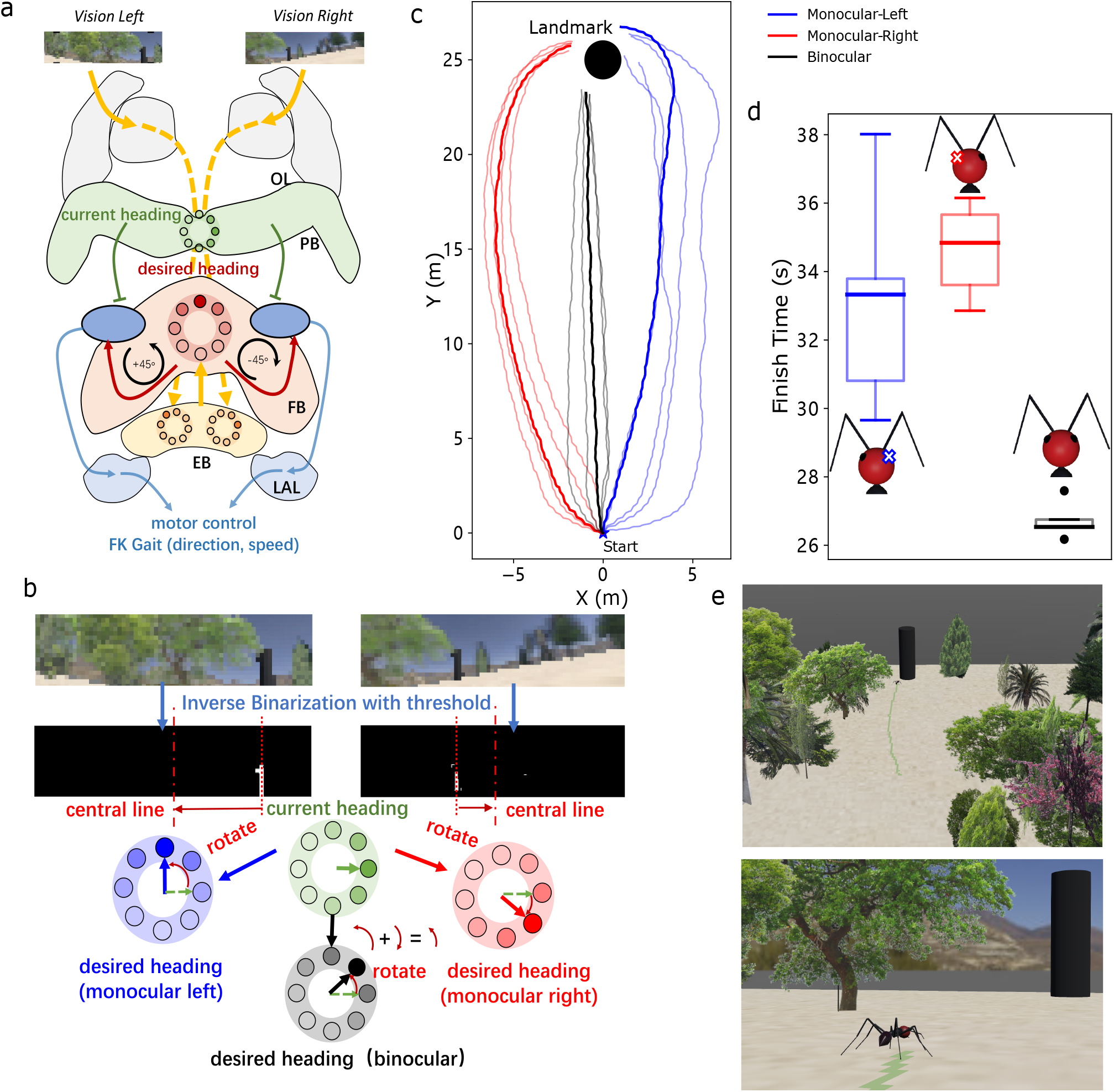
Visual beacons with monocular and binocular vision. (a) Schematic diagram of the visual beacons model overlapped with the insect brain regions. (b) Illustration of the generating desired heading from the visual inputs with monocular and binocular visual input. (c) Trajectories of the visual beacon using left, right, and binocular vision. (d) Time taken by the robot to reach the position near the landmark. (e) Snapshots of the simulated ant robot and environment during visual beacon navigation.

### (ii) Panoramic visual compass

In addition to binocular vision, insects are believed to use panoramic views when navigating [63–65]. Computational models also leverage panoramic images to extract useful information for solving visual navigation tasks, particularly in the frequency domain [29, 66, 67]. Here, we demonstrate how processing panoramic views in the frequency domain can aid the agent in sensing direction, known as the visual compass [65, 68]. The methods used to process the panoramic view here are the same as those in [29, 67], which could also serve as the foundation for insect visual route following behaviour (see Figure 5a). The initially extracted phase encoding in the frequency domain is stored as the desired heading direction. When the simulated 3D world is manually rotated, the received panoramic image and the phase encoding rotate accordingly. The difference between the stored phase and the current phase is output to the steering circuit, enabling the agent to turn and continuously maintain its initial heading. To test the performance of the visual compass-based head tracking, we rotated the 3D world in two modes: incremental rotation and jump rotation (sudden and fast). As shown in Figure 5c and supplementary Movie S8-9, the simulated ant efficiently tracked its stored initial heading in both rotation modes, consistently orienting towards the red ball in the 3D visual world.

**Figure 5.**
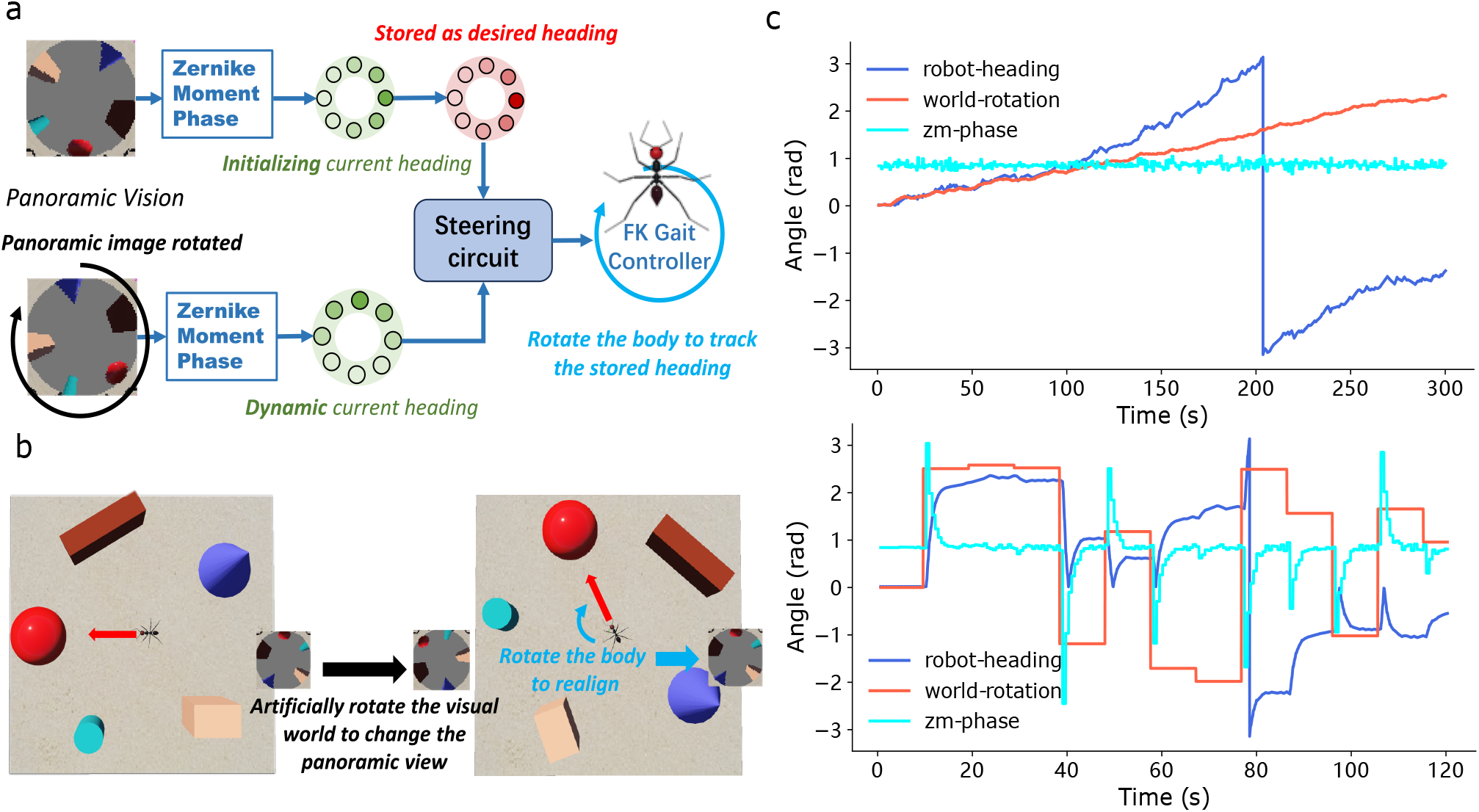
Visual compass using panoramic view. (a) Schematic diagram of the visual compass model. (b) Screenshot of the rotating simulated world from Webots. The rotation mode of the visual scene has two types: incremental (top of (c)) and jump (bottom of (c)). (c) The heading direction of the robot, the visual scene, and the phase of the Zernike moment are plotted when tracking the stored direction by the Zernike moment based visual compass.

### (e) Olfactory-guided manoeuvres

Olfaction represents another crucial sensory domain guiding a diverse array of navigational behaviours [13, 69–71]. Different olfactory landscapes lead to varied navigation behaviours [72]. To showcase how the proposed platform facilitates olfactory navigation simulation under environments with different spatial-temporal features, we implemented odour trail following behaviour under a spatially stable distribution [73] and odour plume tracking behaviours in a turbulent olfactory environment [13].

#### (i) Odour trail following using active sensing

Odour trail following [43] observed in carpenter ants (*Camponotus pennsylvanicus*) was realised to demonstrate how embodiment benefits the design of control strategies. We compared the trail following performance of agents with and without moving their antennae during odour tracking by measuring the time taken to navigate from the same start point to the end of the pheromone trail (finish time as shown in Figure 6d). Intriguingly, we found that agents with moving antennae significantly outperformed those with fixed antennae (Figure 6c,d) when tracking spatially wider trails, although their performance on trails with normal width was similar. This suggests that dynamic antennae movement enhances the adaptability of control strategies to handle a wide range of environmental conditions, as demonstrated in [43]. This finding underscores how embodiment facilitates the design of effective control systems [2].

**Figure 6.**
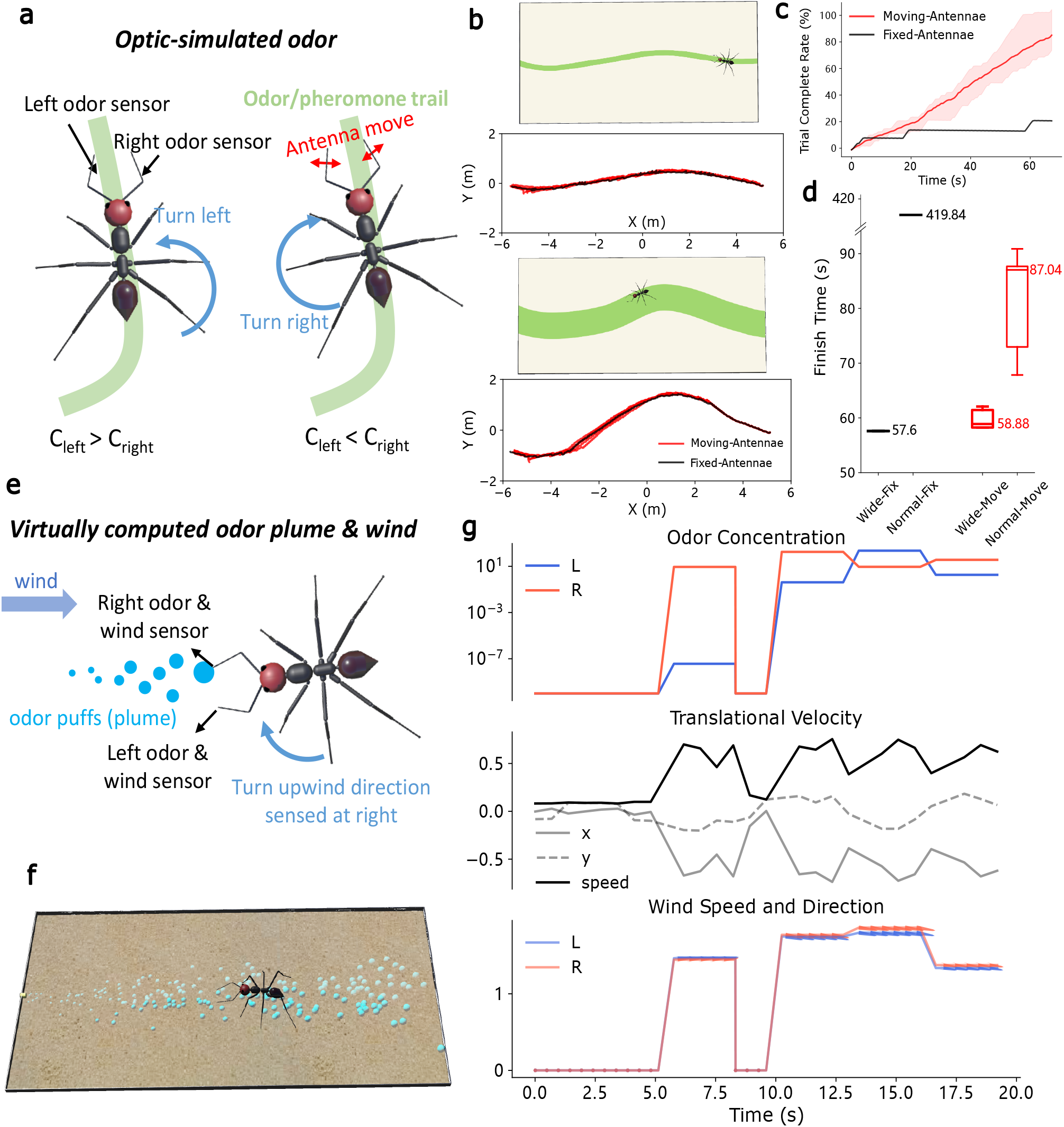
Olfactory-motor closed-loop control. (a)-(d) depict odour trail following, while panels (e)-(g) illustrate odour plume tracking. (a) Schematic diagram illustrating the control strategy of the trail following behaviour. (b) Bird’s-eye screenshots of the experiments and the corresponding trajectories of the agent with different widths of the odour trail. (c) Comparison of the completion rate between fixed-antenna and moving-antenna configurations. (d) Box plot of the finish time (n=5). (e) Model of plume tracking. (f) Screenshot of the experiments in the Webots 3D scene. (g) Perceived odour concentration (top), wind (bottom), and the moving velocity of the ant robot (middle) during one experiment.

In addition, I2Bot simplifies body modification. For example, supplementary Video12 demonstrates a preliminary test where ant robots with varied antenna lengths, but identical control algorithms, exhibited different performances in odour trail following. This together with the results presented in the visual beacon simulations (Figure 4) further highlights the impact of body morphology and physical dynamics on the development of control algorithms and neural circuitry. It underscores that intelligence develops through the intricate interaction between body, brain, and environment[1].

#### (ii) odour plume tracking

To investigate sensory-motor control in turbulent odour environments, we developed a simplified control strategy inspired by behaviours observed in walking flies [13]. While this behaviour has been extensively modeled in previous studies [17, 74], we adapted a conceptual model to validate the proposed tool’s capacity to implement dynamic olfactory navigation behaviours. As illustrated in Figure 6e, once the odour is sensed (i.e., the perceived odour concentration exceeds the threshold), the agent orients upwind (aligning with the wind direction detected via the virtual mechanosensory signal). In the absence of odour, the agent performs random rotations in place, simulating scanning/searching manoeuvres. Despite discontinuous olfactory and mechanosensory cues in this turbulent environment, agents successfully navigate to the odour source using this simple strategy (see Figure 6g and Supplementary Video13).As the first attempt to implement dynamic odour plume tracking in a simulated robot with realistic ant morphology and physical constraints, this case study may provide useful foundational work for future investigations into embodied olfactory-motor control.

## 3. Materials and Methods

I2Bot is build in the popular open-source robot simulator Webots [36]. The controllers of the demos presented in this paper were programmed in Python 3.9 (compatible for newer version) with packages such as *numpy, cv2, ikpy*, etc. Note that aside from Python, Webots supports multiple programming languages like Java, C/C++ and Matlab scripts. All the source code related to this project is open-sourced at Github https://github.com/XuelongSun/I2Bot under the MIT license.

### (a) Robot design

The morphology of a *Cataglyphis fortis* desert ant was estimated from images provided [75] (see Supplemental Fig.S1). To simulate the ant body in 3D, we use Webots embedded geometries *sphere, cone* and *capsule* to model the body parts as that in [39]. As shown in Supplemental Fig.S1, the length of the hind leg is larger, making it distinctive as the characteristics of desert ants’ leg morphology. Note that the size of the ant is scaled 100 times larger for the ease of the physics simulation within Webots.

Binocular Visual sensors are placed at the position of eyes while the abstracted panoramic visual sensor is placed at the top of the ant robot’s head to obtain a more ideal field of view. Olfactory sensors are mounted on the end of the antenna to mimic that of animals [46] and make it easy for robotics studies [47]. Eight tactile sensors are mounted on the tip of each leg and each antenna which can detect three dimensional contact force. Each joint (40 in total) has a torque sensor that could be dynamically accessed during locomotion (see Figure 1). This torque feedback could be very useful in designing bio-plausible locomotion controllers like that in [48, 51].

### (b) Environment construction

For visual environments, all visible objects, such as basic geometric shapes and trees used in the presented cases, are available in Webots. Vision is constructed using the camera model embedded in Webots. Additionally, simulated visual environments from previous studies on insect navigation can be easily imported into the Webots 3D environment via the *Mesh* shape or *Cadshape* node. See Supplemental Fig.S2 for examples of visual scenes used in [27, 29, 67, 76]. This feature allows researchers to directly obtain visual stimuli from the desired world without needing to configure a camera model, facilitating the comparison of different visual navigation models that share the same visual input.

For olfactory environments, there are two types of simulation: 1) for the odour trail, it is optically simulated as the specific texture of the floor. This texture can be drawn manually thus allows great flexibility. 2) for the odour plume, it is simulated by the filament-based model [77] and its Python implementation called *pompy*. This model mimics both short- and long-timescale features of odour plume evolving in a turbulent flow and has been widely used in olfactory related studies involving insect navigation [69, 78] and robot odour source localization [79].

### (c) Models

#### (i) Forward kinematics

To simply the gait control, we keep the ThCx and TiTa joint angles constant and denote the angles of the left three joints-ThCz, CTr and FTi to be *α, β* and *γ* respectively. These values are determined by the hip swing (*S*_*l*_ with unit *rad*), lift swing (*S*_*h*_ with unit *rad*) and the moving direction (−1 for backwards and 1 for forwards). To realise yaw control of the robot body, we introduced rotation configuration denoted as *θ*. Table 2 lists the formulas to calculate the base range of the ThCx joint *R*.

**Table 2.** The calculation of ThCx joint range *R* for yaw-control of forward kinematics gait.

|  | left legs (FL, ML and HL) | right legs (FR, MR and HR) |
| --- | --- | --- |
| $\theta \geq 0$ | $S_l d(2\theta - 1)$ | $S_l d$ |
| $\theta < 0$ | $S_l d(2\theta + 1)$ | $-S_l d$ |

As different gait types share the same mechanisms, here we just describe how to use FK to generate a tripod gait, other gait could be generated with alternative leg coordination sequence. For the first half phase of a leg during the swing stage, the angles of the three joints are calculated by:

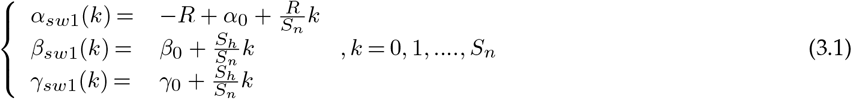

where *α*_0_, *β*_0_, *γ*_0_ is the initial angle of ThCx, CTr and FTi joint respectively. *S*_*n*_ is the total time steps to lift the leg to its **11** highest position determined by the lift swing *S*_*h*_. Similarly, in the next half phase, the angles are calculated by:

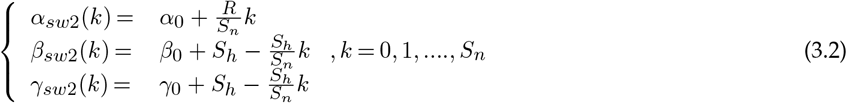

In this phase, the leg is moved to the frontmost position (*α* reaches its highest value) and contacts the ground again (*β* and *γ* are set to its initial value).

For the stance phase, *β*(*k*) and *γ*(*k*) are kept constant (as the leg does not lift) while *α*(*k*) is reversed (moves from the front of the body to its back):

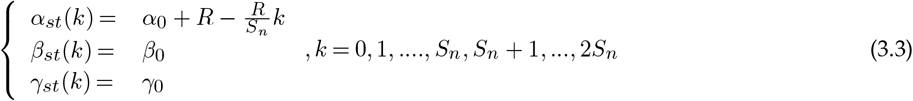

For a tripod gait, the angle of ThCx joint of FR, ML and HL leg should be *α*_*sw*1_ -> *α*_*sw*2_ -> *α*_*st*_ while that of the other three legs (i.e., FL, MR and HR) should be *α*_*st*_ -> *α*_*sw*1_ -> *α*_*sw*2_. The angles of CTr (*β*) and FTi (*γ*) joint follow the same rule.

#### (ii) Inverse kinematics

Unlike the forward kinematics wherein the joint angles are computed directly given the leg movement parameters (i.e., hip swing *S*_*l*_, lift swing *S*_*h*_, etc.), when applying inverse kinematics, we first calculate the positions of the leg tip at a certain time step *k* in swing and stance state, and then utilize the inverse kinematics (with python package-*ikpy*) to compute the joint angles. The legs of the ant robot were defined by URDF files and can be read by the *ikpy* package. Specifically, the step length and step height is denoted as *L* and *H* respectively, then for leg in the swing stage, the joint angles should be:

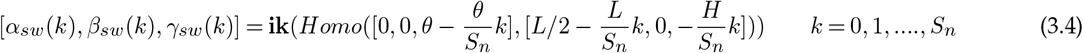

Where *Homo*(**rotation, translation**) calculates the homogeneous matrix given the rotation and translation input and **ik**() denotes the IK function which receives the homogeneous matrix of the leg tip as parameters and returns the calculated joint angles (This function is provided as an API in the *ikpy* package, and the specific calculations are not detailed here, as they follow standard inverse kinematics processes). *θ* is the rotation configuration. For the stance leg:

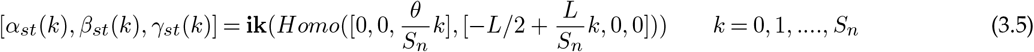

Thus, to generate a tripod gait, the angle of the ThCx joint of FR, ML and HL leg should be *α*_*sw*_ -> *α*_*st*_ while that of the other three legs (i.e., FL, MR and HR) should be *α*_*st*_ -> *α*_*sw*_. The angles ofthe CTr (*β*) and the FTi (*γ*) joints follow the same rule.

#### (iii) Path integration

Path integration requires heading direction and velocity information, this is provided by the **Webots** *Supervisor* in the current implementation. One can use other modalities like vision (e.g. optic flow) to obtain this. The model of path integration is adapted from [28, 29], the output of the summed CPU1 (PFN) neurons are fed into the FK-based gait controller as following:

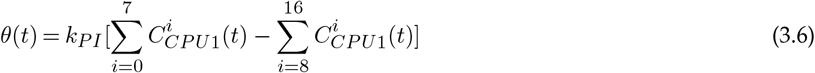

where 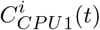 is the membrane potential of the *i*^*th*^ CPU1 neuron at time *t*. Similar with that in [28], the difference between the summed activation of left and right CPU1 neurons modifies the rotation configuration of the gait control (see Table 2 for specific calculation) and thus guides the robot’s turning decision via the motor scale *k*_*PI*_. To tune the walking speed, step length *S*_*l*_ is computed by:

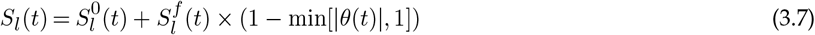

where the basis step length 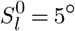 and the scale factor 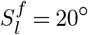 are held constant. Thus, the step length is confined in the range of [5°, 25°]. These parameters are selected empirically based on the ant robot’s performance in the designed simulation. For running multiple trials (results presented in Figure 3d), we set the start location to be fixed at [0, 0] while the initial heading varies in range [0, 2*π*].

#### (iv) Vision **12**

Vision is provided through the embedded camera sensor in Webots. For panoramic view simulating, the projection mode of the camera is set to *spherical* while for monocular and binocular it is set to be *planar*. The parameters of the camera such as resolution, field of view and focal length can be customized in Webots easily through the *scene tree*.

- **Visual beaconing**: The resolution of the left and right eye camera is 74 × 19 (i.e., the image width *W*_*vb*_ = 74 and the image height *H*_*vb*_ = 19) with the horizontal and vertical field of view to be 2*rad* and 0.5*rad* respectively. The motor command is calculated by the ’copy-and-shift’ mechanism proposed in [17, 29]. The input image will be first binaries with a given threshold (empirically selected in the range [10, 25] given that the brightness of the image is in the range [0, 255]). Then the value of *shift*_*vb*_ is computed by:
- 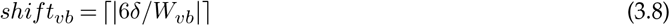

Where *δ* is the different between the horizontal image center and the center of the landmark area:

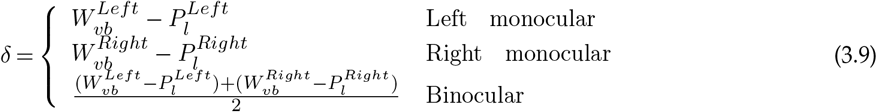

Where *P*_*l*_ is the averaged horizontal position of the dark landmark in the left and right retina (i.e., image pixel coordinates). The desired heading is then calculated through the *copy-and-shift* mechanism wherein the current heading is copied and shifted by *shift* amount. Then the turning value *θ*(*t*) is calculated by the steering circuit [17, 28] as that in Equation 3.6 but with a different motor scale of visual beacons *k*_*vb*_. The ThCx joint range *R* (Table 2) and the hip swing *S*_*l*_(*t*) (Equation 3.7) of the FK gait controller then could be calculated to affect the locomotion. For the multiple trials simulation, the ant robot starts from the same point but with different initial headings. Specifically, for agents using the left eye view uniformly sample from the range [0, *π/*2], whereas for agents using the right eye view sample uniformly from the [*π/*2, *π*]. For binocular agents, they sample from the range [*π/*4, 3*π/*4].
- **Visual compass**: The resolution of the artificial panoramic vision is 72 × 72. The direction of the view is extracted by the phrase information of the Zernike moment [29, 67] coefficient (with order *n* = 7 and repetition *m* = 1, *Φ*_7,1_(*t*)). This Zernike phase is compared with the initially stored phase (*Φ*_7,1_(0)) and then the difference determines the sign of turning parameter *θ*(*t*) and the hip swing:

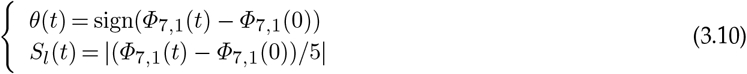

#### (v) Olfactory

- **Odour trial following**: As described above, the odour trail is simulated by the texture of the floor, thus the concentration is detected by the *DistanceSensor* (note that the field type should be set to be “infra-red”) whose value is modified by a reflection factor depending on the color, roughness and occlusion properties of the object. The detected odour concentration of the odour (returned value of the *DistanceSensor*) from the left and right antenna at time *t* are denoted as *O*^*l*^(*t*) and *O*^*r*^(*t*), the difference between the left and right sensed odour concentration is denoted as *O*_*d*_(*t*) = *O*^*l*^(*t*) − *O*^*r*^(*t*). Table 3 shows how the sensed odour concentration determines the locomotion control (i.e., determines the hip swing and rotation of the FK gait control). For multiple trial experiments, all the agents started from the same position [5, 0] with identical initial heading (*π*).
- **Odour plume tracking**: In this simulation, the agent not only detects the odour concentration (*O*^*l*^ and *O*^*r*^) but also the wind direction 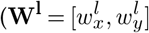 and 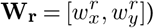. These values are computed virtually based on the filament-based model [77]. When there is no odour sensed, the agent conducts a random searching around the spot (set *θ* to be -1 or 1 randomly and *S*_*l*_ to be a small value like 5°). While the odour is sensed (i.e., the concentration value exceeds 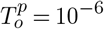), the agent compares its left and right odour concentration and turns to the up-wind direction sensed with higher odour concentration. Specially, the desired turning orientation is computed by *m*(*t*) = tan^*−*1^(*w*_*y*_*/w*_*x*_). Then the step length and rotation are set to be:

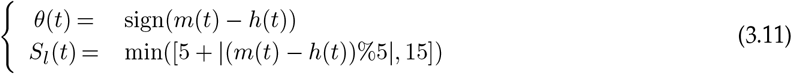

where *h*(*t*) is the current heading of the ant robot. Note that the agent will conduct 1 gait loop using the parameters defined in the above equation and then going forward (*S*_*l*_ = 15, *θ* = 0) for 2 gait loop.

**Table 3.** odour trail following. The threshold *Td* = 5 *To* = 900 are set empirically according to the lookup table of the *DistanceSensor* used. **1***Kd* = 400 is a constant scale factor.

| sensory state | odour sensed |  | no odour |
| --- | --- | --- | --- |
| | $O_d < T_d$ | $O_d \leq T_d$ | $O^l < T_o$ & $O^r < T_o$ |
| $\theta$ (degree) | 0 | $O_d/k_d$ | -1 or 1 (random) |
| $S_l$ (degree) | 8 | 4 | 3 |

## 4. Discussion

Computer simulations have proven extremely valuable in neuroethology studies of insect vision (e.g. [80, 81]), brain function (e.g. [28, 29]), and locomotion (e.g. [53, 82]) allowing researchers to verify and adapt their hypotheses efficiently and effectively. Yet, as animal behaviour arises from the interaction of each of the above plus information from internal and external cues, new tools are required that integrate as many of these elements as possible. Moreover, for these tools to be adopted widely across this highly multidisciplinary community they should be robust, easy to install, and easy to use.

As a step towards this goal, we presented I2Bot, an open-source tool for studying the mechanisms of multimodal and embodied insect navigation. We have provided a series of case studies that demonstrate the usability and flexibility of the proposed platform, which has the potential to accelerate research in this fast moving research field. Compared to other simulation tools (see Table 4), Hector [83] and NeuroMechFly [33] primarily focus on locomotion. Additionally, I2Bot provides a flexible visual and olfactory world construction and is simple to extend with other tools. For example, CompoundRay [84] could be integrated with the camera model to generate more realistic insect visual inputs, and make use of more realistic 3D environments (e.g., [85–87]). Regarding the physics engine, both the Open Dynamic Engine (ODE) and MuJoCo are popular in the robotics field, each with its own pros and cons for physics simulation (for detailed comparison, see [88, 89]).

**Table 4.**
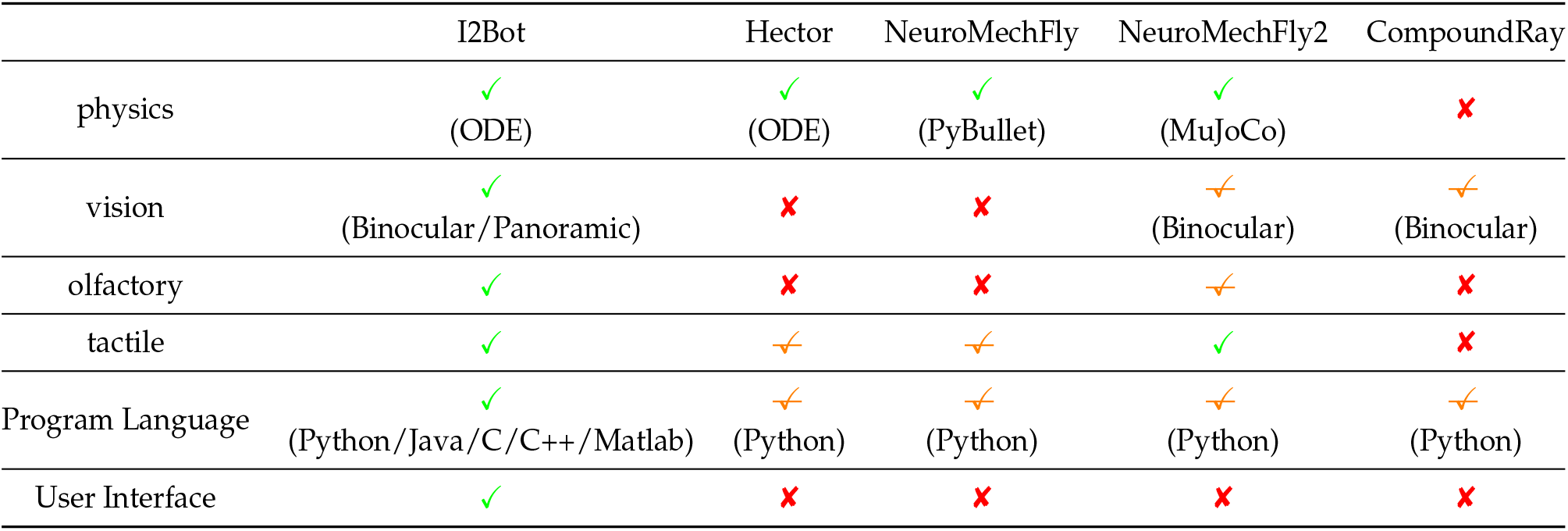
Comparison between different simulation tools.

### (a) Reinforcement Learning and Swarm Intelligence

I2Bot can also be used as a (deep) reinforcement learning (RL/DRL) tool [90–92], which is becoming a more common tool for insect neuroscience [34, 78, 93] and insect-inspired robotics [53, 94], but requires efficient simulation run the large number of trials involved in the optimisation process. The proposed tool could provide an efficient way to conduct this kind of study. The sensory and body kinematic states of the robot can construct the *State Space*, while the motor commands sent to all available joints form the *Action Space*. Our robot acts as an RL *agent* could dynamically interact with the controllable *environment. Reward* is scenario dependent, for example, in the visual navigation context, the distance between the agent and the desired spot could be used to form the *reward* while in the odour plume tracking case, *reward* could be determined by the sensed odour concentration. The proposed I2Bot platform offers a high variety of interaction forms that interest researchers from biology and robotics, facilitating research into the underlying mechanisms of intelligent behaviours and the training and testing of RL-based algorithms.

I2Bot is also suitable for simulating multi-agent systems, making it ideal for swarm intelligence and robotics studies [95, 96]. Similar to previous studies using Webots for swarm robotics simulation [97–99], I2Bot can easily be applied to investigate similar scenarios by adding multiple ant robots (see supplementary Video12 for an example of multiple agents simulation). This approach can lead to embodied swarm intelligence, as the body morphology and physical dynamics are integrated into this platform.

### (b) Roadmap for a community hub: simulation tools in insect neuroethology

The intention of this study was to demonstrate the opportunities that modern computer simulation tools offer to the field of insect neuroethology, taking the classic desert ant navigation problem as a representative use case. Yet, for I2Bot’s maximum value to be realised will require adoption by the community to create a virtuous cycle of usage driving tool-development and vice versa. We outline a roadmap of developments that could take place to enhance the tool to make it usable by the broad community of insect neuroethologists in Table 5. Such extensions will allow researchers to continuously enhance the platform’s capabilities and apply it to a broader range of studies in insect navigation and robotics. Our hope is to enable community hub to drive] ever faster research cycles as has been realised through tool sharing in field such as machine-learning (such as FastAI[100] and MLlib[101]), robotics (like ROS [102]) and neuroscience (such as FlyWire [103] and insectBrainDB [104]). To this end, we extend an open invitation to the neuroethology community to become active contributors to this open-source project (https://github.com/XuelongSun/I2Bot).

**Table 5.** Roadmap for developing a community hub of simulation tools in insect neuroethology **14**.

|  | Now<br>(I2Bot currently can offer) | Next<br>(implementable immediately) | Future<br>(long-term featured goals) |
| --- | --- | --- | --- |
| **Sensory** | Vision (Binocular and Panoramic);<br>Olfactory (Airborne and chemical);<br>Force sensing (joint torque and tactile) | More bio-realistic vision simulation (e.g., embedded CompoundRay[84]);<br>More precise tactile sensing with touch sensor arrays;<br>Compound odour sensing | Multimodal and real-time sensory |
| **Brain Models** | Central Complex steering circuit[28];<br>Heading direction using ring attractor;<br>Central Complex copy-and-shift[17] | Mushroom bodies model for visual navigation[27];<br>Models of the insect motor centre:lateral accessory lob(LAL)[105];<br>Central complex multiple cues integration[17], etc. | Bio-plausible neural models of insect brain |
| **Locomotion** | First Body model (desert ant);<br>Robotics gait control | Biomimetic locomotor control(e.g., NeuroWalkNet[51]);<br>Body models of other walking insects (e.g.,stick insect [82]);<br>Embedded other body model framework[106], etc. | Agility of walking, flying and jumping |
| **Environment** | 3D Visual scenes;<br>Odour plume with wind;<br>Stable odour trails;<br>Uneven terrain and walls | Visual scenes reconstructed from real world;<br>Polarise light simulation;<br>Magnetic simulation;<br>simulating multiple odours simultaneously, etc. | Realistic 3D and interactive environment |

## A. Supplementary files

Supplementary files are listed as below:

Fig.S1. The design of I2Bot’s morphology is based on real desert ant’s measurement.

Fig.S2. Examples of importing previous used visual 3D world [14, 27, 29, 67, 76] into I2Bot environment.

Movie S1. Gait control using forward kinematics.

Movie S2. Gait control on uneven terrain.

Movie S3. Wall climbing using adhesion.

Movie S4. Simulation of path integration.

Movie S5. Visual beacons using monocular left eye.

Movie S6. Visual beacons using monocular right eye.

Movie S7. Visual beacons using binocular vision.

Movie S8. Visual compass (jump mode).

Movie S9. visual compass (incremental mode).

Movie S10. odour trail following (normal trial).

Movie S11. odour trail following (wide trail).

Movie S12. odour trail following with varied antenna length.

Movie S13. odour plume tracking.

### Ethics

This work did not require ethical approval from a human subject or animal welfare committee.

### Data Accessibility

All the source codes, model of the robot and environments are open-sourced via Github https://github.com/XuelongSun/I2Bot under the MIT license.

### Authors’ Contributions

X. Sun., M. Mangan. and S. Yue. conceived the idea and designed the experiments. X. Sun. developed the tool and conducted the experiments. All authors contributed equally to the writing of the manuscript.

### Funding

This research has received funding from the National Natural Science Foundation of China under the Grant No 62206066 and No 12031003.

