## Supplementary figures and images for "I2Bot: an open-source tool for multi-modal and embodied simulation of insect navigation"

### Fig,S1

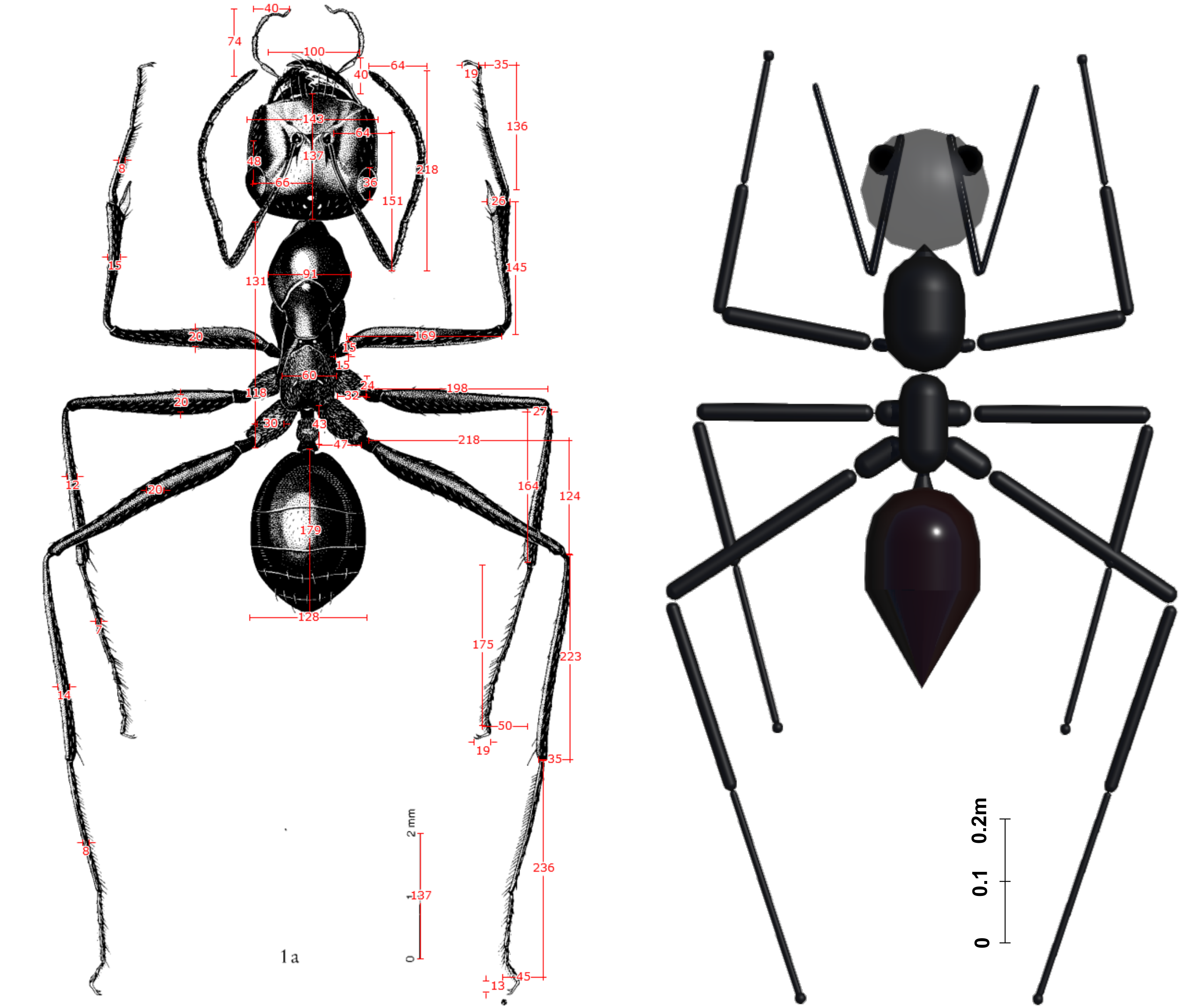

### Fig,S2.

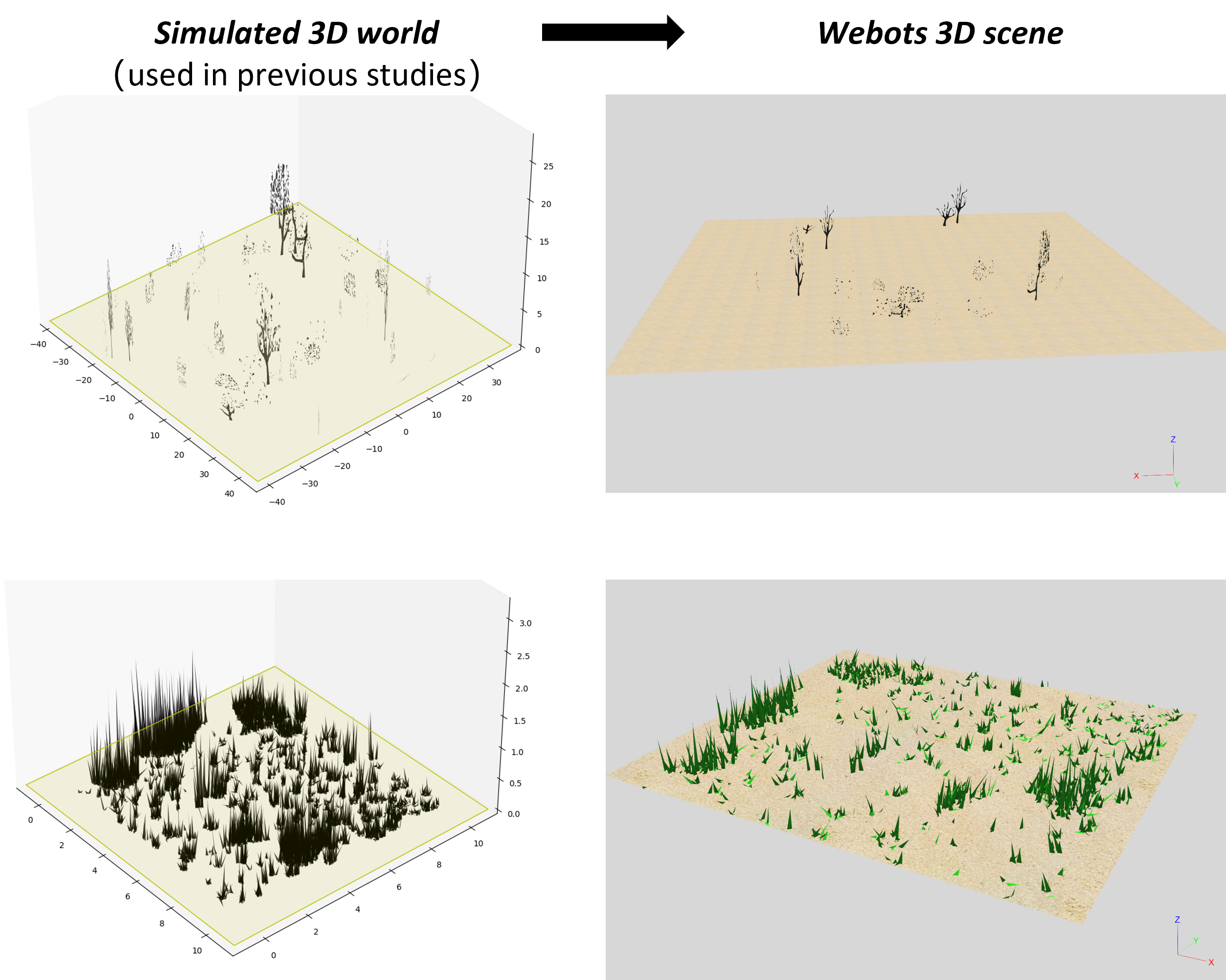
